# Loss of GdpP function in *Staphylococcus aureus* confers β-lactam-specific antibiotic tolerance and promotes invasive infection

**DOI:** 10.64898/2026.07.27.740951

**Authors:** Vedangi D. Hayatnagarkar, Stefano G. Giulieri, Raymond Poon, Swagata Bose, Joshua B. Parsons, Steven Y. C. Tong, Vance G. Fowler, Benjamin P. Howden, Som S. Chatterjee

## Abstract

The emergence of antibiotic tolerance in *Staphylococcus aureus* reduces antibiotic efficacy by allowing bacterial survival despite prolonged antibiotic exposure, the molecular basis of which remains poorly understood. Moreover, the phenotypic indistinguishability of tolerant isolates in antimicrobial susceptibility testing impedes effective diagnosis and therapy. Increased concentration of the second-messenger, cyclic-di-AMP (CDA), has recently been implicated in tolerance to β-lactams as well as other cell-wall-reactive antibiotics. Using the ScanLag assay, Tolerance-Disk test, and traditional methodologies and employing isogenic mutagenized strains, we demonstrate that loss of GdpP function, a phosphodiesterase that hydrolyzes CDA, confers tolerance specifically to β-lactam antibiotics independent of their class. The extent of β-lactam tolerance correlated directly with the intracellular CDA concentration and inversely with the inhibition of bacterial cell-wall synthesis. Δ*gdpP* mutants caused higher mortality than wild-type strains in the *Galleria mellonella* infection model upon β-lactam treatment, suggesting GdpP-mediated tolerance could lead to β-lactam treatment failure. Large-scale within-host evolution analysis demonstrated that MRSA and MSSA strains isolated from patients acquire GdpP loss-of-function mutations during invasive infections but not during nasal carriage. Overall, this study highlights the clinical relevance of *gdpP* mutations, frequently selected in persistent *S. aureus* infections, as key mediators that could promote treatment failure due to β-lactam tolerance.

**Importance:** Although GdpP loss-of-function mutations have been associated with antibiotic tolerance in MRSA and MSSA, whether they directly mediate tolerance, the antibiotic specificity of this phenotype, and how they contribute to infections remain unresolved. Our study shows that an enhanced level of cyclic-di-AMP (CDA), which results from GdpP loss-of-function mutations, promotes tolerance specifically to the β-lactam class of antibiotics in *Staphylococcus aureus*. The mechanism by which such tolerance is conferred is independent of the classical β-lactam resistance genes, *mecA* and *blaZ*. This report also provides evidence that GdpP loss-of-function mutations may promote persistent invasive infections in patients and points to potential future steps to re-sensitize CDA-dependent β-lactam-tolerant *S. aureus* strains.

## Introduction

The Gram-positive bacterium, *Staphylococcus aureus*, is both a common human colonizer and an opportunistic pathogen responsible for a wide spectrum of diseases, including nosocomial and community-associated infections, such as abscesses, as well as invasive infections, such as bacteremia and osteomyelitis (1, 2). It is the leading cause of bacteremia-associated global mortality (3). It is a formidable pathogen due to its potential to evade antimicrobial interventions through multifaceted antibiotic resistance and tolerance mechanisms, coupled with pronounced virulence (4–6).

β-lactam antibiotics, such as antistaphylococcal penicillins, cefalosporins including cefazolin, are highly efficacious and safe for treating *S. aureus* infections (7, 8), especially those caused by methicillin-susceptible strains (MSSA). However, these β-lactams, including next-generation ones (NGBs), are not effective against Methicillin-resistant (MRSA) strains due to resistance conferred by the low-affinity Penicillin-Binding Protein, PBP2a. Effective alternatives to NGBs, such as highly advanced cephalosporins (ceftaroline and ceftabibrole), vancomycin, daptomycin, and linezolid or their combinations, are currently used to treat MRSA infections (9, 10).

Antibiotic tolerance has recently received significant attention in the field for its ability to cause treatment failure. Tolerance is a transient physiological adaptation to antibiotic-induced stress that enables bacteria to better survive the lethal action of antibiotics. Bacteria can tolerate high antibiotic pressure through adapted physiological changes rather than genetic mutations in resistance genes (11). Not only is the mechanism of antibiotic tolerance distinct from that of antibiotic resistance, but they are also distinct phenotypically (12). For example, while resistance to antibiotics is always associated with an increase in drug MIC, antibiotic tolerance is not. As a result, identification of tolerant bacterial strains is difficult, as they remain phenotypically susceptible to the tested antibiotic. Moreover, mechanisms that promote antibiotic tolerance can be multifarious in nature, such as slower growth and prolonged lag phase, etc., which remain poorly characterized, unlike the mechanisms that promote antibiotic resistance (11, 13), further contributing to the problem of diagnosing antibiotic-tolerant bacterial strains.

Cyclic-di-AMP (CDA), a recently discovered bacterial second messenger, has been postulated to drive antibiotic tolerance of various cell wall-reactive antibiotics, including vancomycin and daptomycin, among Gram-positive bacterial pathogens (14–18). In particular, loss-of-function mutations in GdpP, a <u>p</u>hosphodiesterase that cleaves CDA to its linear form, 5’-phosphadenylyl-adenosine (pApA) (19), are increasingly detected in clinical strains of *S. aureus* that are non-susceptible to β-lactam antibiotics (20–22). These mutations in GdpP lead to an accumulation of CDA in bacteria and allow enhanced survival when challenged with β-lactam antibiotics through mechanisms that haven’t been clearly identified (14, 15, 23). Besides antibiotic tolerance, CDA also plays crucial roles in regulating bacterial cell size, cell wall stress, and ion homeostasis (24–28). It has recently been reported that increased CDA levels in bacteria lower cellular turgor pressure, prevent bacterial cell lysis, and reinforce the peptidoglycan network (29). In addition to its broad role in bacterial physiology, CDA has also been implicated in playing critical roles in bacterial pathogenesis (30). Previous studies have reported an association between loss-of-GdpP-function mutations with decreased bacterial virulence and attenuated innate immune response during *S. aureus* infections (31, 32). Moreover, recent clinical studies have identified *S. aureus* isolates with loss-of-GdpP-function mutations from patients during episodes of infection, underscoring the clinical importance of CDA in staphylococcal pathogenesis (33–35).

In this study, we aimed to determine the antibiotic classes against which loss of GdpP function mediates tolerance. For this purpose, we used a clinical *S. aureus* USA300 strain in which we previously identified GdpP loss-of-function mutations in passaging experiments. We deleted *gdpP* from the Wild-type *S. aureus* USA300 strain (Wt) and from its *mecA* and *blaZ* excised variant (Wtex) to create Δ*gdpP* mutants (36, 37). Antibiotic tolerance of the isogenic strain pairs was studied using a real-time optical plate-scanning assay (ScanLag) and a tolerance disk test (TD). A wide range of antibiotic classes, comprising several cell wall targeting antibiotics (e.g., vancomycin, daptomycin, bacitracin, and a diverse range of β-lactam antibiotics spanning multiple generations, including ceftaroline), antibiotics that target bacterial protein synthesis (e.g., erythromycin, chloramphenicol, and tetracycline), and antibiotics targeting DNA replication and transcription (e.g., ciprofloxacin and rifampicin, respectively), were tested. Our findings indicated that loss of GdpP function confers tolerance specifically to β-lactam drugs, regardless of class, and that the extent of this tolerance is directly correlated with the intracellular abundance of CDA in *S. aureus*. We also observed that the displayed β-lactam tolerance of the Δ*gdpP* strains depended on bacterial cell wall synthesis. *In vivo* infection studies carried out in *Galleria mellonella* demonstrated that β-lactam tolerance due to the lack of *gdpP* increased worm mortality upon β-lactam treatment compared with their isogenic parental strains. Within-host evolution genomic analysis was performed on a large cohort of clinical MRSA and MSSA isolates from bacteremia and nasal carriage infections. Our results suggested that GdpP loss-of-function mutations occur during persistent invasive infections in humans. Collectively, this report emphasizes the role of GdpP’s loss of function in β-lactam tolerance and its clinical importance during *S. aureus* infections.

## Materials and methods

### Bacterial strains

*S. aureus* strains were cultured in tryptic soy broth (TSB) media (BD Biosciences, USA), with aeration at 180 rpm or on tryptic soy agar (TSA) (BD Biosciences) plates and incubated at 37°C. Complemented strains were constructed by introducing the constitutively expressing *pTX*Δ plasmids into the recipient strains via phage transduction. Strains containing *pTX*Δ plasmids were cultured in TSB media or on TSA plates, supplemented with 12.5 mg/L tetracycline. All strains and plasmids used in this study are listed in **Tables S1 and S2**. Clinically derived USA300 background strain (Wt), USA300 with excision of *mecA* and *blaZ* (the classical mediators of β-lactam resistance in *S. aureus*), and their isogenic *gdpP* deletion mutant strains (Wt Δ*gdpP* and Wtex Δ*gdpP*) were used predominantly for this study.

### MIC assay

The MIC assay was performed using the broth microdilution method as previously described (38). Briefly, 1 × 10^5^ colony-forming units (CFU) of bacteria were incubated in 0.2 mL of cation-adjusted Mueller-Hinton broth (BD Biosciences) containing increasing concentrations of antibiotics. The concentration range used for nafcillin, oxacillin, cefazolin, ceftriaxone, vancomycin, daptomycin, chloramphenicol, and tetracycline was 0.25-256 mg/L. For ceftaroline and imipenem, the range was 0.03-4 mg/L; for erythromycin, it was 0.06-8 mg/L; for bacitracin, 4-4096 mg/L; and for ciprofloxacin and rifampicin, 0.003-4 mg/L. MICs were defined as the lowest concentration that inhibited bacterial growth after incubation at 37 °C for 48 h. The MIC assay was performed twice to ensure reproducibility. The antibiotics used in the study are listed in **Table 1** and **Table S3**.

**Table 1:** Antibiotics used in the study and MICs (mg/mL) of Wtex and Wtex Δ*gdpP*. Note: Clinical & Laboratory Standards Institute (CLSI) breakpoints are based on CLSI (M100-ED35), 35th edition (S: susceptible and R: resistant).

| Antibiotics | Classification and function | MIC (µg/mL) |  | CLSI breakpoints |
| --- | --- | --- | --- | --- |
|  |  | Wtex | ΔgdpP |  |
| β lactams (Prevent cell wall synthesis by inactivating PBPs) |  |  |  |  |
| Nafcillin | Next-generation β-lactam | 1 | 1 | S |
| Oxacillin | Next-generation β-lactam | 2 | 2 | S |
| Cefazolin | Cephalosporin (1<sup>st</sup> generation) | 1 | 2 | S |
| Ceftriaxone | Cephalosporin (3<sup>rd</sup> Generation) | 8 | 8 | S |
| Ceftaroline | Cephalosporin (5<sup>th</sup> generation) | 0.5 | 0.5 | S |
| Imipenem | Carbapenem (1<sup>st</sup> generation) | 0.06 | 0.125 | S |
| Antibiotics targeting cell envelope |  |  |  |  |
| Vancomycin | Glycopeptide (Binds to D-ala-D-ala and prevent peptidoglycan cross-linking) | 1 | 1 | S |
| Bacitracin | Polypeptide antibiotic (Prevents the final dephosphorylation step in the phospholipid carrier cycle) | 32 | 32 | S |
| Daptomycin | Cyclic lipopeptide (causes membrane depolarization and ion-leaks) | 1 | 1 | S |
| Antibiotics targeting protein synthesis |  |  |  |  |
| Erythromycin | binds to ribosomal 50S subunit | 0.5 | 0.5 | S |
| Chloramphenicol | binds to ribosomal 50S subunit and inhibits peptidyl transferase | 16 | 16 | R |
| Tetracycline | binds to ribosomal 30S subunit | 1 | 1 | S |
| Antibiotics targeting DNA replication and transcription |  |  |  |  |
| Ciprofloxacin | Fluoroquinolone (Inhibits bacterial DNA gyrase) | 1 | 1 | R |
| Rifampicin | Rifamycin (Inhibits bacterial RNA polymerase) | 0.015 | 0.015 | S |

### TD test

The tolerance disk (TD) test was performed as described previously with some modifications (39, 40). Briefly, overnight *S. aureus* cultures were adjusted to an OD600 of 0.1 in TSB, and 200 μL of the bacterial suspension was plated onto TSA plates. Complemented strains containing *pTX*Δ plasmids were plated on TSA plates with 12.5 mg/L tetracycline. A 6 mm paper disk (Becton Dickinson) was placed on TSA plates containing plated bacteria, 10 μL of antibiotic solution was added to the disk, and the plates were incubated at 37°C for 24 h. The absolute amount of each antibiotic was equal to twice its MIC. For the combinatorial treatment, to achieve the required absolute amounts on the disk, antibiotic solutions at twice the usual concentration were mixed 1:1 (v/v), and 10 μL of the mixture was added to the disk. **Table S3** describes the concentration, dilutions, absolute amounts, and solvent used for each antibiotic. After 24 h of incubation, the zone of inhibition was measured to note the inner 50% of it at a later stage, and 4 mg of glucose, i.e., 10 μL of 40% D-glucose (Sigma), was added onto the same paper disk. For *mecA*-positive strains, 10 μL of 20% glucose (2 mg) was added to the disk, except for tests with nafcillin, in which 4 mg of glucose was added. Plates were incubated at 37°C for an additional 48 h. Following the incubation, tolerant colonies in the inner 50% of the zone of inhibition were enumerated using the microscope (Leica EZ4W, Evos XL Core), plotted, and analyzed using GraphPad Prism. TD tests were performed in triplicate, and the experiment was repeated twice to ensure reproducibility.

### ScanLag assay

The colony appearance time of different *S. aureus* strains on TSA plates containing antibiotics was measured with a flatbed scanner and the ScanLag program software package, following a previously described protocol with some modifications (41). Briefly, *S. aureus* strains were grown overnight in 4 mL of TSB at 37 °C and 180 rpm. The overnight cultures were diluted with PBS to a bacterial suspension with an OD600 of 5, then serially diluted to 10^-6^. 100 µL of the 10^-6^ diluted bacterial suspension was plated onto TSA plates with and without antibiotics. Antibiotic concentrations were optimized so that 100-250 colonies would be on the plate when the above dilution was plated. Autoclaved black fabric was sandwiched between the TSA plates’ top and the bottom, ensuring the fabric does not touch the agar. The plates were placed bottom side down on an EPSON Perfection V200 flatbed scanner, and the scanner was placed in a 37°C incubator. Images were taken at every 30-min interval, starting after 240-min of bacterial plating. The resulting images were analyzed using the ScanLag analysis package written in MATLAB. Data were exported from MATLAB and analyzed using Excel and GraphPad Prism. The data in the figure are a representative experiment of 3 biological replicates. Statistical analysis was performed with the 3 biological replicates.

### Quantification of Cyclic-di-AMP

Intracellular CDA concentration was measured using a riboswitch as previously described (42). Briefly, *S. aureus* strains were freshly transformed with the plasmid *pTX*Δ [*bsuO* P6-4], and three individual colonies were chosen for each strain. The chosen riboswitch clones were grown in culture tubes with 4 mL of TSB containing 12.5 µg/mL Tetracycline overnight at 37°C at 180 rpm. Bacterial culture with an OD600 of 500 was collected and centrifuged for 5 min at 6000 g. The cell pellet was washed with 1 mL of PBS and resuspended in 100 µL of TSB containing 12.5 µg/mL Tetracycline and 200 µM DFHBI-1T. The resuspended bacterial cultures were incubated at 37°C for 1 h in the dark. After incubation, 40 µL of labeled bacterial culture was diluted into 1 mL of PBS. 200 µL of the diluted cells was put in a round-bottom 96-well plate, 2 wells per sample. The plate was placed into a Guava easyCyte Flow cytometer (Luminex, USA). 30000 events were collected using the blue laser (488 nm excitation) and the green channel (512-530 nm emission). The data were analyzed using InCyte software (GuavaSoft, version 4.0). The mean fluorescent intensity from the GFP channel was calculated for each sample and plotted using GraphPad Prism.

### Galleria mellonella infection model

Larvae of *Galleria mellonella* (greater wax moth) were procured from Vanderhorst Wholesale, Inc. (Ohio, USA). The *in-vivo* infection and worm survival assays were performed using a previously reported protocol (43) with slight modifications. Briefly, larvae were kept on wood chips in darkness at 25 °C during the sixth instar stage. Larvae weighing 200–300 mg were selected and used within a week of receipt. For each experimental condition, 20 healthy larvae were incubated in sterile 9-cm Petri dishes lined with filter paper and acclimatized for 2 h at 37°C before infection. 3 × 10⁵ CFU bacterial suspensions were used for infecting each larva. Inoculation was done by injecting 10 µL suspension of the bacteria into the hemocoel via the last left proleg using 25 µL syringes (Hamilton). After infection, larvae were incubated at 37°C for 168 h and were observed every day for survival. Each experimental group consisted of a total of 20 larvae. For negative controls, larvae were infected with heat-killed wild-type strains. A heat-killed bacterial suspension was prepared by incubating 3 × 10⁵ CFU at 65°C for 20 minutes, then tested by plating 100 µL on TSA plates to confirm growth inactivation, with cell morphology retained and assessed microscopically. For antibiotic-treated infection assays, 0.5 µg/mL nafcillin was injected into larvae 2 h post-infection. An additional control group treated with 0.5 µg/mL nafcillin alone was included to assess potential drug toxicity, and no toxicity was observed. Infection with heat-killed bacteria and nafcillin alone served as negative and positive controls, respectively. Infection with both *mecA* +ve and -ve strains and their Δ*gdpP* mutants was tested without or with nafcillin treatment. Kaplan–Meier larval survival curves were plotted. Dead worms were identified by failure to respond to gentle touch at the posterior tip with sterile forceps. The experiment was done twice to ensure reproducibility. Statistical analysis was performed using the log-rank (Mantel-Cox) test using GraphPad Prism.

### Invasive infection studies and computational analysis

A genomic meta-analysis was performed of published within-host evolution studies of *S. aureus* sterile-site infection (bacteremia, bone infection, soft-tissue infection) and nasal carriage. Infection/colonization episodes were extracted from a large-scale within-host evolution study published in 2022 (33), supplemented with 3 bacteremia studies (34, 44, 45) and one nasal carriage study (46). A variant-calling approach was applied, which is specifically designed for within-host evolution studies and is described in (33). Briefly, all reads were mapped to an internal reference (either a complete genome or draft assembly generated using Shovill, v1.1.0 [https://github.com/tseemann/shovill]) using Snippy, v4.6.0 (https://github.com/tseemann/snippy); variants from internal reference reads and variants in regions of low coverage were removed using scripts available at https://github.com/stefanogg/staph_adaptation_paper. Mutated proteins were clustered using CD-HIT, v4.8.1 (47), and homologs in the reference genome USA300_FPR3757 (Accession number: GCF_000013465.1) were identified using Blastp. Within-host phylogenies were reconstructed as in (35) with outgroup genomes selected from all other episodes based on genetic distance calculated using Mash, v2.3 (48). The ancestral state of *gdpP* was inferred based on the outgroup genomes.

### Statistical Analysis

A One-way analysis of variance (ANOVA) or a two-tailed unpaired *t-test* or the log-rank (Mantel-Cox) test was used to analyze statistical significance using GraphPad Prism software (version 8.0, La Jolla, CA, USA). Significance levels were represented by **\*** (*P* ≤ 0.05), **\*\*** (*P* ≤ 0.01), **\*\*** (*P* ≤ 0.001), **** (*P* ≤ 0.0001) and ns for non-significance. The significance of protein variants was inferred by modelling independent mutation counts (i.e., one protein variant per infection episode) for each USA300_FPR3757 homolog using Poisson regression; to infer the significance of the GdpP variants enrichement we used likelihood ratio to compare the actual mutation counts with the null hypothesis, represented by a uniform mutation count for each gene. The statistical analysis was performed separately for invasive infection and nasal carriage episodes. To account for the small dataset and the short-term within-host evolution, the dN/dS ratio calculation was adjusted by adding pseudo-counts (+1) to the non-synonymous (substitutions and truncations) and synonymous counts (49).

## Results

### GdpP-loss-of-function-mediated antibiotic tolerance is specific to the β-lactam class of drugs

We have previously reported that the loss of GdpP function in *S. aureus* can lead to β-lactam tolerance (14). Our results were in agreement with another study that identified similar loss of GdpP function mutations in an oxacillin-passaged laboratory strain (17). The same study also reported that loss of GdpP function can cause vancomycin tolerance in *S. aureus* (17). Moreover, *gdpP* mutations were detected in patients with persistent bacteremia who underwent daptomycin therapy (45). Thus, whether GdpP loss-of-function mediated tolerance is restricted to β-lactams or can it also mediate tolerance to other antibiotic classes, including other cell wall reactive drugs such as vancomycin and daptomycin, remained unclear. To determine this, we studied the ability of a *gdpP* deletion mutant to produce tolerance to a broad range of antibiotics. This included β-lactams of various types such as next-generation β-lactams (nafcillin and oxacillin), cephalosporins (cefazolin, ceftriaxone, and ceftaroline), and carbapenem (imipenem). In addition, other antibiotics targeting the bacterial cell envelope (vancomycin, bacitracin, and daptomycin), DNA replication (ciprofloxacin), transcription (rifampicin), and the translational machinery (erythromycin, chloramphenicol, and tetracycline) were also tested in this study. The modes of action and targets of antibiotics in each category are illustrated in **Fig. 1**.

**Figure 1:**
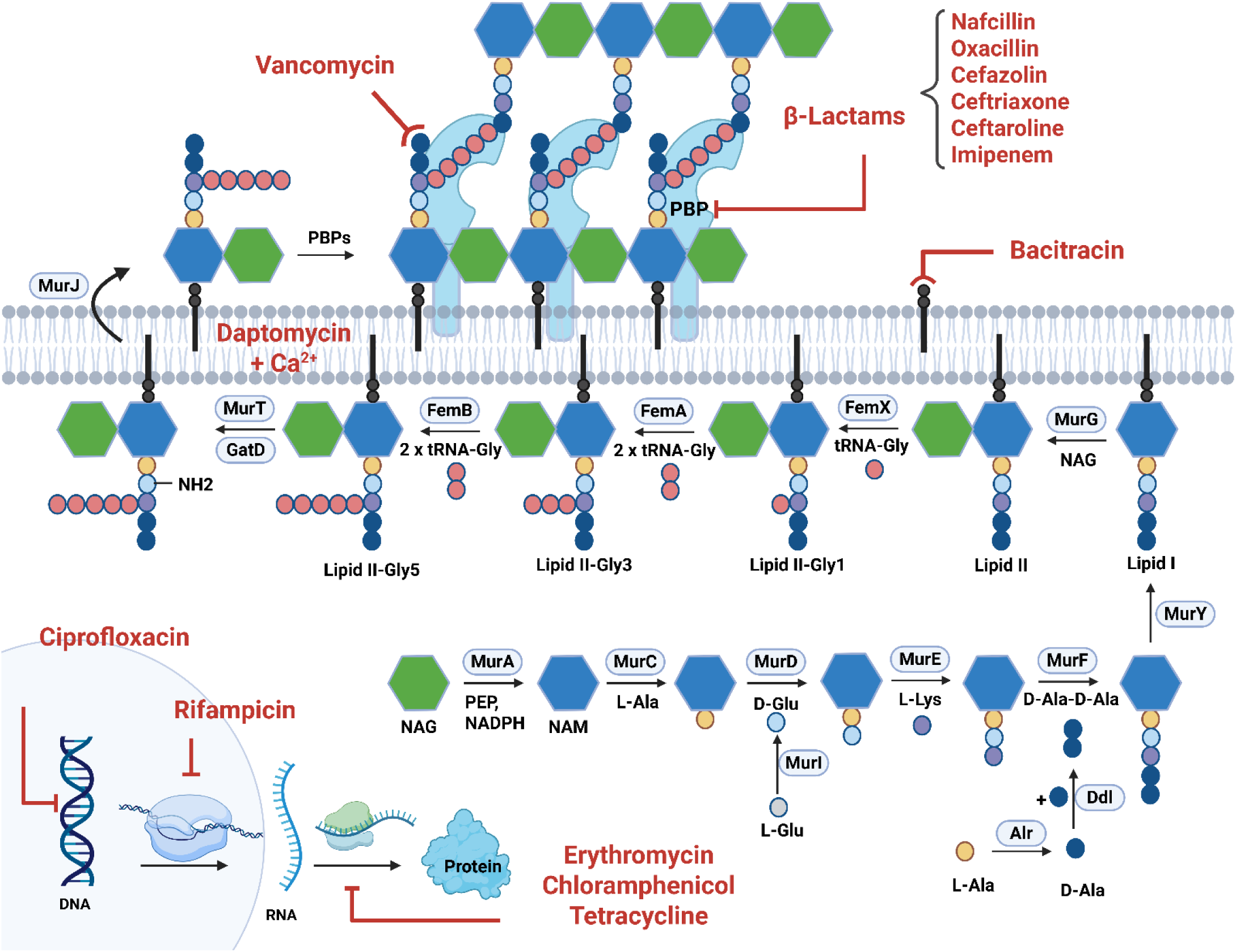
Schematic representation of the staphylococcal cell wall synthesis pathway, the antibiotics used in this study, and their targets. The cell wall synthesis pathway is modified from (74). The figure was created with BioRender.com.

The MICs of the listed antibiotics in **Table 1** were tested for the Wtex and its Δ*gdpP* mutant strain. Both strains displayed similar MICs for all antibiotics except cefazolin and imipenem, which showed a two-fold increase in the Δ*gdpP* mutant but remained within the susceptible range as per CLSI guidelines (**Table 1**). These results indicated that loss of GdpP function mediates antibiotic tolerance but not resistance. The tolerance disk test (TD test) was performed to identify specific classes of antibiotics that confer Δ*gdpP*-mediated tolerance (**Table S3**). This enables identification of antibiotic tolerance by evaluating bacterial survival on agar plates (39). Results of the TD test indicated that the Δ*gdpP* strain produced tolerance exclusively to the β-lactam class of drugs, irrespective of their types (**Fig. 2**). Unlike β-lactams, none of the other antibiotics tested produced Δ*gdpP*-mediated tolerance in our study (**Fig. 3**). Rifampicin and tetracycline treatment caused non-significant differences in tolerant colonies between the Wtex and its isogenic Δ*gdpP* strains, likely due to the appearance of random mutations following treatment with these antibiotics (50, 51).

**Figure 2:**
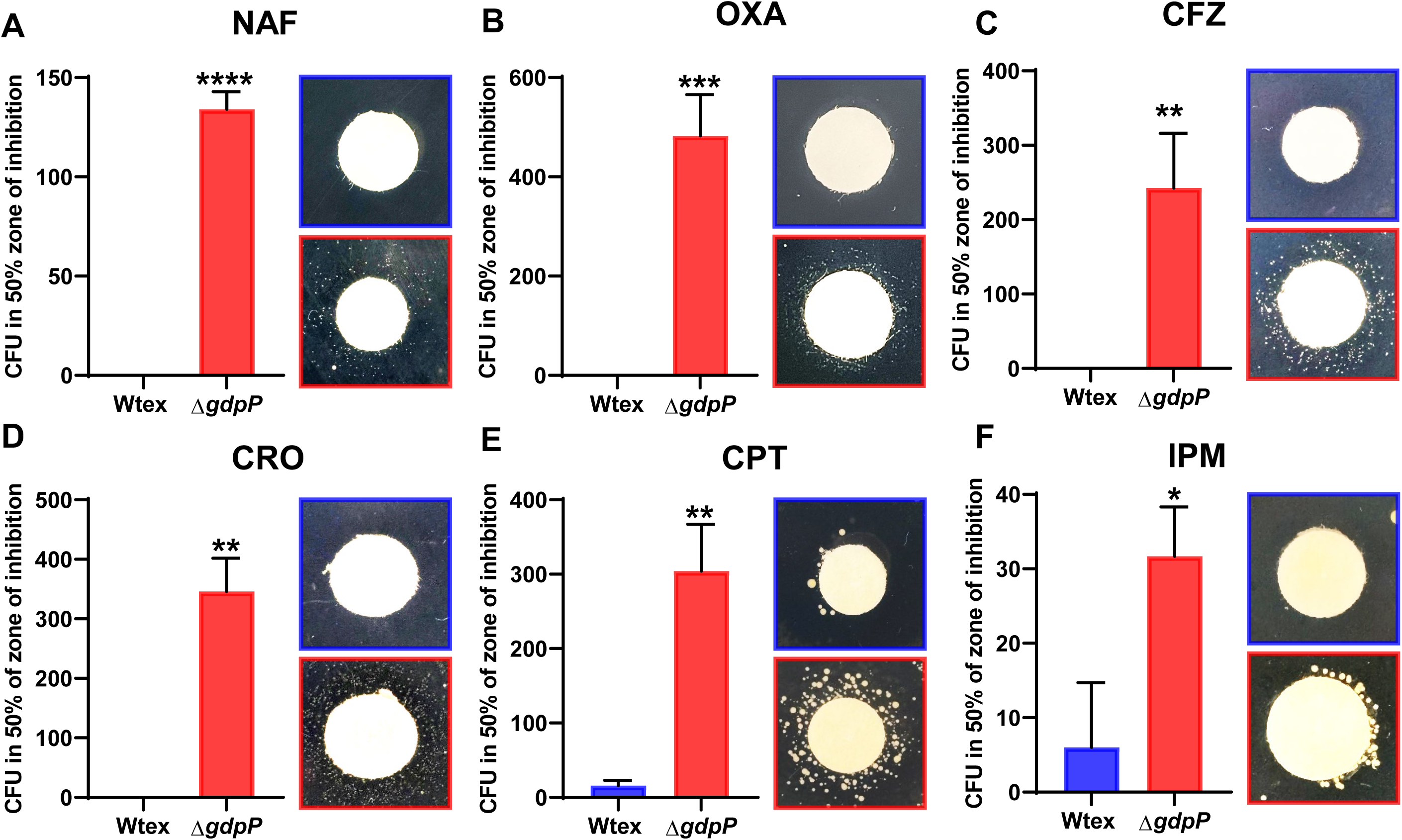
Loss of GdpP function mediates tolerance to β-Lactam drugs. Tolerance Disc (TD) test was performed using the β-lactam antibiotics nafcillin (A), oxacillin (B), cefazolin (C), ceftriaxone (D), ceftaroline (E), and imipenem (F). The concentrations used for antibiotics are listed in **Table S2**. Bacterial CFU in the inner 50% zone of inhibition were enumerated and plotted. The Δ*gdpP* strain (red) showed significantly greater tolerance than its isogenic Wt strain (blue) in all cases. Statistical significance was evaluated using an unpaired two-tailed t-test. * represents *P* ≤ 0.05, ** represents *P* ≤ 0.01, *** represents *P* ≤ 0.001 and **** represents *P* ≤ 0.0001.

**Figure 3:**
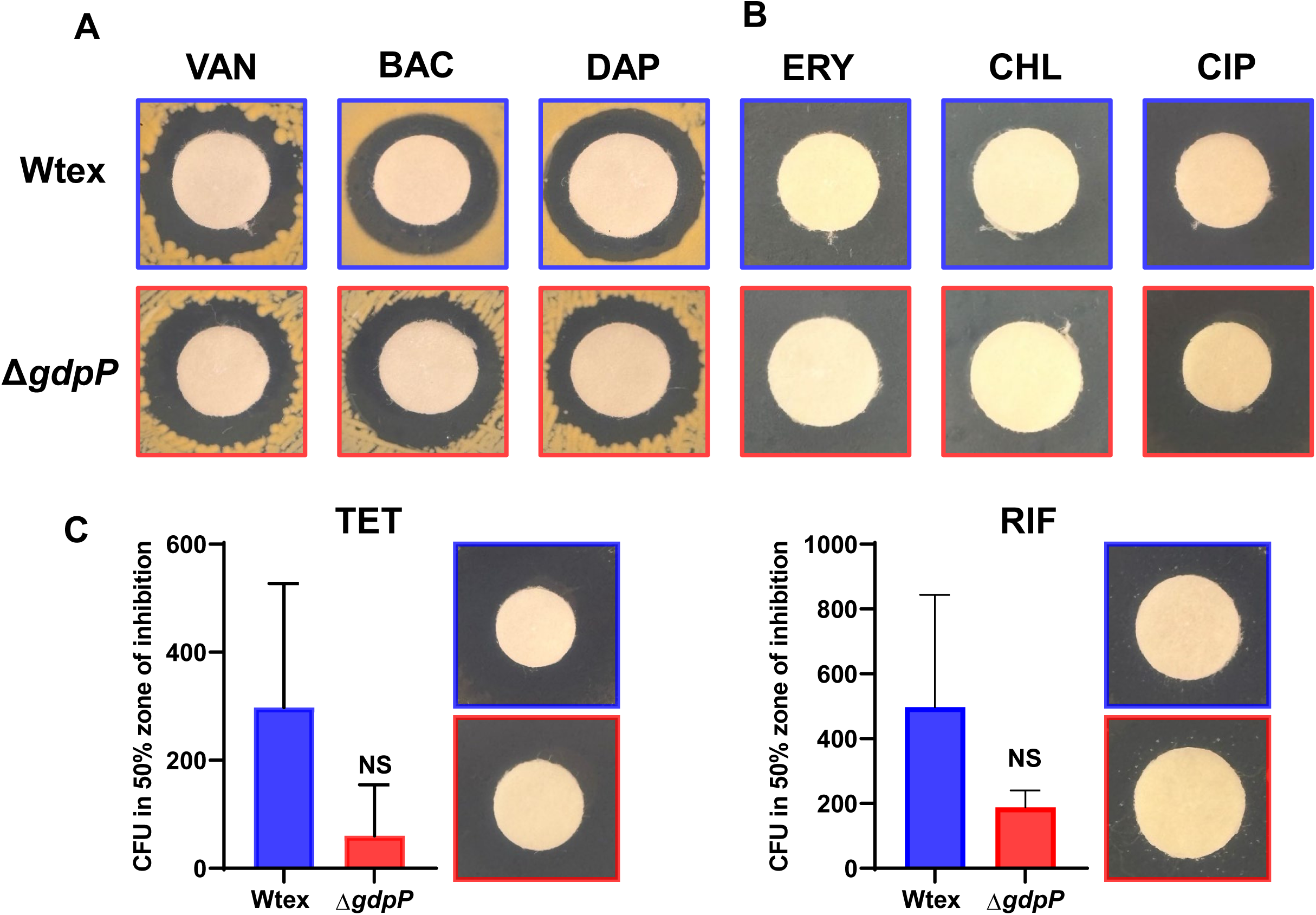
**Loss of GdpP function does not mediate tolerance to non-β-Lactam antibiotics**. Tolerance Disc (TD) test was performed using cell-envelope reactive non-β-lactam antibiotics (vancomycin, bacitracin and daptomycin), antibiotics targeting protein synthesis (erythromycin, chloramphenicol, and tetracycline), and antibiotics targeting DNA replication and transcription (ciprofloxacin and rifampicin, respectively). The concentrations used for antibiotics are listed in **Table S2**. (A) No tolerant colonies were observed using antibiotics, vancomycin, daptomycin, and bacitracin, erythromycin, chloramphenicol and ciprofloxacin. (B) Microscopic colonies appeared on plates treated with tetracycline and rifampicin. The Bacterial CFU in the inner 50% zone of inhibition were not significantly different between Δ*gdpP* (red) and its isogenic Wt strain (blue). Statistical significance was evaluated using an unpaired two-tailed t-test. NS represents non-significant.

To determine if Δ*gdpP* also produces antibiotic tolerance that is specific to the β-lactams in strains that possess *mecA* and *blaZ* (i.e., MRSA strains), we performed the TD test with a Wt USA300 strain and its isogenic Δ*gdpP* mutant. Our results indicated that both nafcillin and ceftaroline conferred a significantly higher level of tolerance due to loss of GdpP, whereas treatment with vancomycin and daptomycin did not lead to tolerance, and the MIC values were also similar for both strains (**Fig. S1**). Thus, Δ*gdpP* in the MRSA background also confers tolerance specifically to β-lactams, similar to that observed in strains lacking *mecA* and *blaZ*.

The β-lactam-specific tolerance of the Δ*gdpP* strain was also determined through an independent assay using ScanLag analysis (**Fig. 4**), which employs automated quantitative measurement of colony appearance on agar plates (41). Our results indicated that in the untreated condition, the Δ*gdpP* colonies appeared slower than its isogenic Wtex strain, likely due to an inherent growth defect, known to be present in Δ*gdpP* strains (40). Furthermore, we observed that although treatment with antibiotics such as nafcillin, vancomycin, daptomycin, or bacitracin was able to delay the appearance of colonies, only in the case of NAF was this delay greater in Wtex compared to the Δ*gdpP*. In the presence of other antibiotics (vancomycin, daptomycin, or bacitracin), the delay time difference (i.e., the difference between no antibiotic vs antibiotic-treated conditions) was greater in the Δ*gdpP* than in Wtex (**Fig. 4**). Collectively, these results further substantiated the fact that Δ*gdpP* mediates tolerance selectively to the β-lactam class of antibiotics.

**Figure 4:**
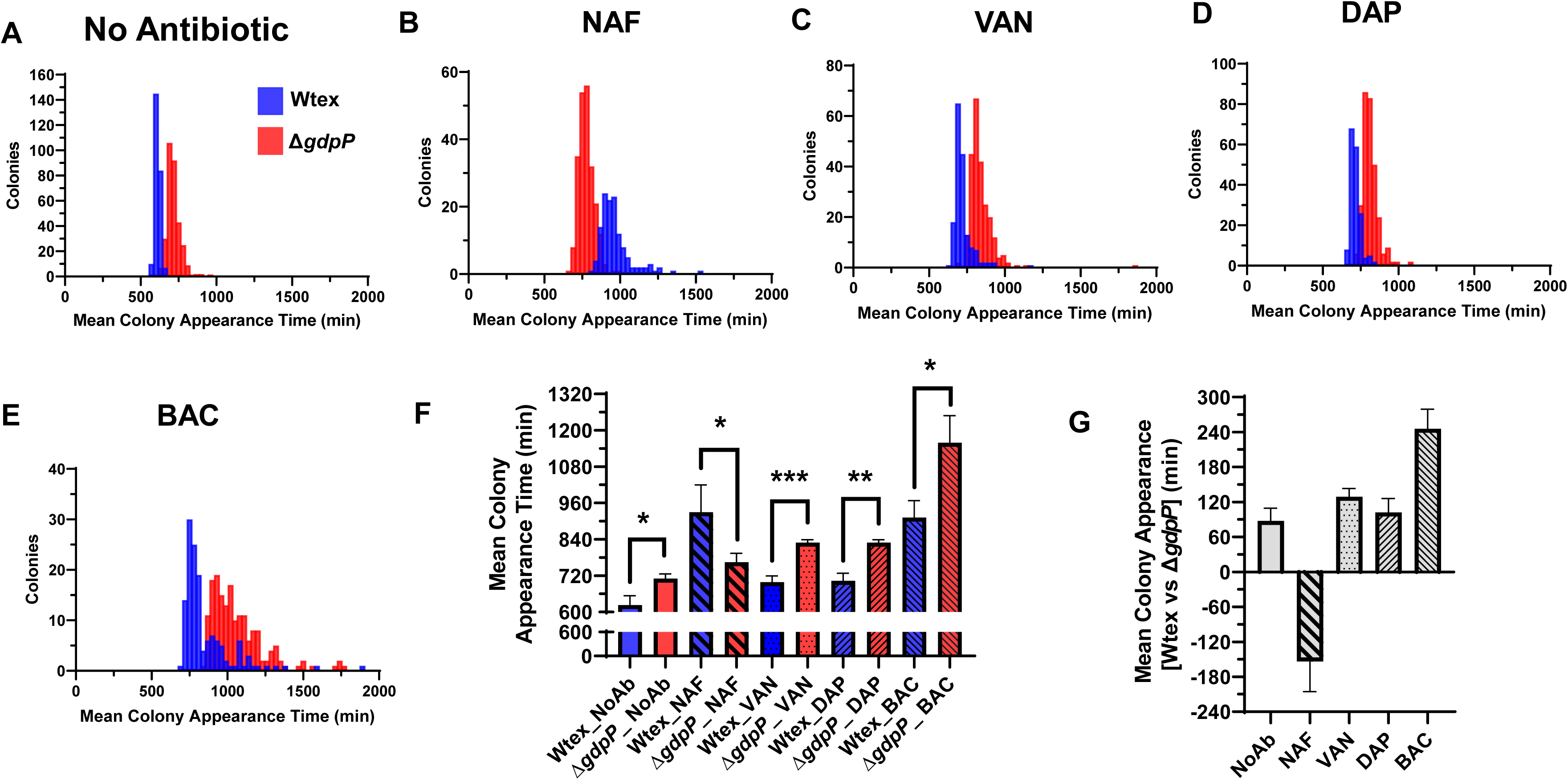
Antibiotic tolerance mediated by GdpP loss-of-function is specific to β-Lactam drugs. (A-E) ScanLag assay showing bacterial colony appearance time on agar plates with or without antibiotics. Colonies for Δ*gdpP* (red) appeared slower than those for its isogenic Wt strain (blue) on plates without antibiotics. The appearance of the Δ*gdpP* colonies occurred faster than that of the Wt strain in the presence of nafcillin due to β-lactam tolerance. This tolerance effect was not detected in similar assays carried out with vancomycin, daptomycin, and bacitracin. (F) Amalgamated analysis of results from biological triplicates of ScanLag assay with no antibiotic (NoAb), NAF (0.125 μg/mL), VAN (0.5 μg/mL), DAP (0.5 μg/mL), BAC (16 μg/mL). The mean colony appearance time is significantly shorter in the Δ*gdpP* mutant than in its isogenic Wtex strain, only in β-Lactam (nafcillin) treated condition. (G) Mean colony appearance time of Δ*gdpP* as compared to its isogenic Wtex strain. Statistical significance was evaluated using an unpaired t-test. * represents *P* ≤ 0.05, and ** represents *P* ≤ 0.01 and *** represents *P* ≤ 0.001.

### Loss of GdpP function-mediated β-lactam tolerance correlates with intracellular CDA levels in a concentration-dependent manner

Since loss-of-function mutations in *gdpP* have been shown to elevate intracellular cyclic-di-AMP (CDA) levels in both laboratory-passaged and clinical strains in our earlier studies (14, 42), we sought to investigate whether β-lactam tolerance observed in the Δ*gdpP* strains correlates with CDA concentrations. To test this, the Δ*gdpP* strain was complemented with Wt *gdpP*, two mutants of *gdpP* (*gdpP*-V609D and *gdpP*-Q642X) having intermediate CDA levels, and the empty vector *pTX*Δ. Intracellular CDA concentrations in these *gdpP*-complemented strains were previously described in our study (14). Tolerance levels were evaluated using a TD test with ceftriaxone treatment and plotted against the corresponding CDA concentrations for each complemented strain. β-lactam tolerance mediated by *gdpP* mutations exhibited a positive correlation with CDA concentration (**Fig. 5**). The correlation curve (**Fig. 5A**) represented a rapid rise in tolerance effect, even with a slight increase in CDA level, and saturation of tolerance effect at higher CDA concentrations. Ceftriaxone was chosen for these studies because it usually produces a high dynamic range needed for the experiment. These results indicated a direct association between CDA levels and β-lactam tolerance mediated by loss of GdpP function.

**Figure 5:**
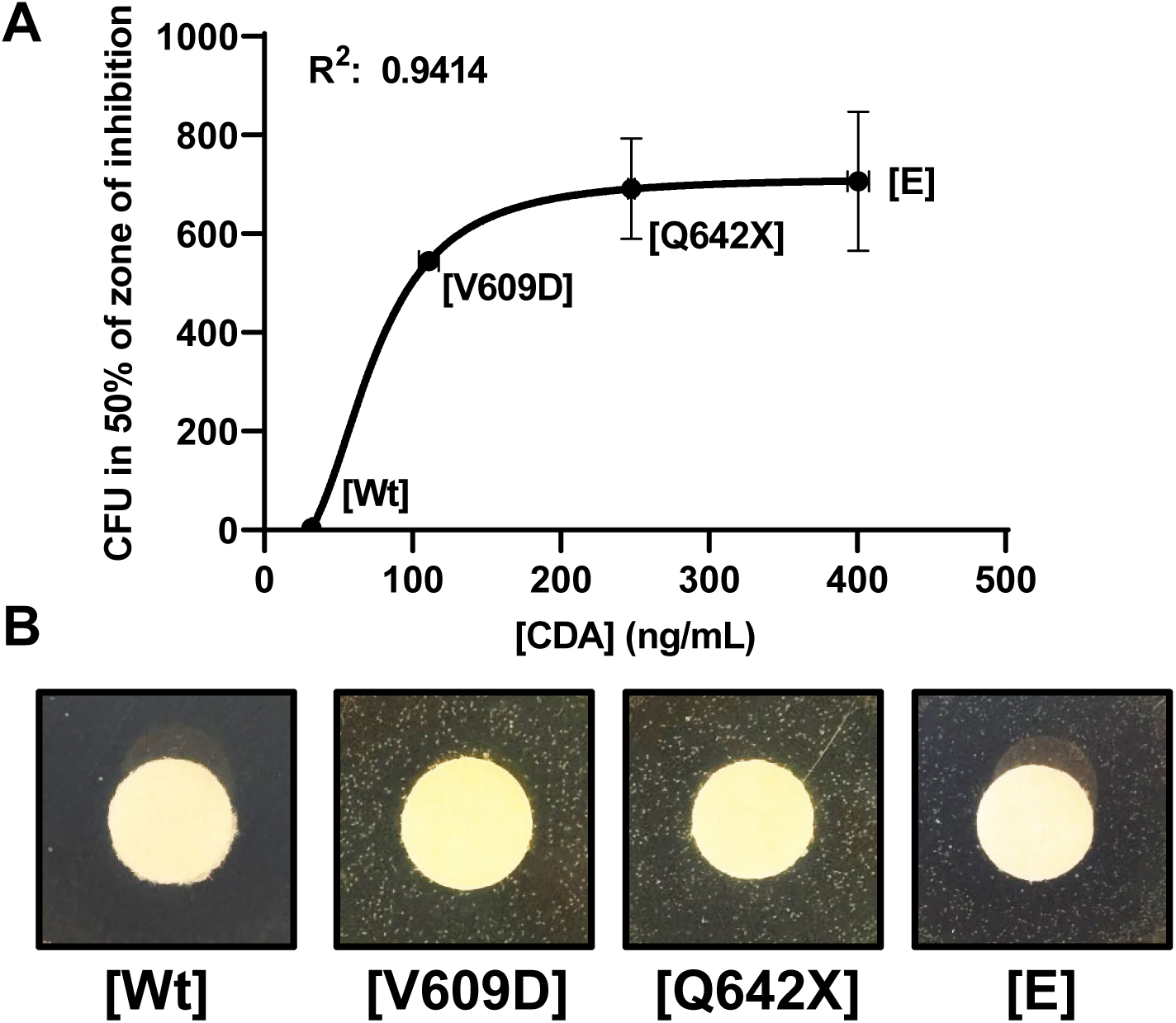
β-Lactam tolerance mediated by GdpP loss-of-function is CDA concentration dependent. Tolerance ability of the Δ*gdpP* strain complemented with Wt *gdpP* [Wt], two mutant *gdpPs* [*gdpP* V609D], [*gdpP* Q642X], and the empty vector [E] was plotted against their respective CDA levels identified previously (14). A high correlation coefficient (R^2^=0.9414) (A) shows that the β-Lactam tolerance (B) of these strains is positively dependent on their intracellular CDA concentrations.

### β-lactam-tolerant Δ*gdpP* colony isolates from TD tests are not resistant mutant variants

Five tolerant Δ*gdpP* colonies were picked from the inner 50% of the zone of inhibition of the TD test with nafcillin. These isolated nafcillin-tolerant colonies were analyzed through a subsequent TD test with β-lactams, nafcillin, cefazolin, and ceftaroline (**Fig**. **6**). The results indicate that all five tolerant Δ*gdpP* colony isolates are significantly more tolerant to β-lactams than Wtex. Similar tolerance levels were observed among the majority of tolerant Δ*gdpP* colony isolates relative to their parental Δ*gdpP* strain. Some variability in tolerance levels seen among the isolated colonies is most likely due to spontaneous mutations. It has been previously shown that increased CDA concentrations in bacteria promote higher mutation rates, enabling them to evolve faster and better survive antibiotic pressure (52). Similar to the parent Δ*gdpP* strain, MIC values of nafcillin, cefazolin, and ceftaroline for the isolated tolerant-Δ*gdpP* colonies were in the susceptible range as per the CLSI guidelines (**Fig.6C**). Intracellular CDA concentrations of the tolerant-Δ*gdpP* isolates were determined using a CDA riboswitch by transforming them with a vector containing a fluorescent biosensor (*bsuO* P6-4) as described previously (42) (**Fig**. **S2**). Intracellular CDA levels of the isolated tolerant-Δ*gdpP* colonies were similar to those of their parent Δ*gdpP* strain, with no significant difference, but were significantly higher than those of Wtex. Collectively, these results demonstrated that nafcillin-tolerant Δ*gdpP* colony isolates are not resistant mutant variants. Slight differences in MICs, which are within the susceptible range, and minor differences in tolerance potential are likely due to some secondary mutations that these colonies acquire during their growth and experimental procedure, as tolerant strains are more prone to acquire secondary mutations (53).

**Figure 6:**
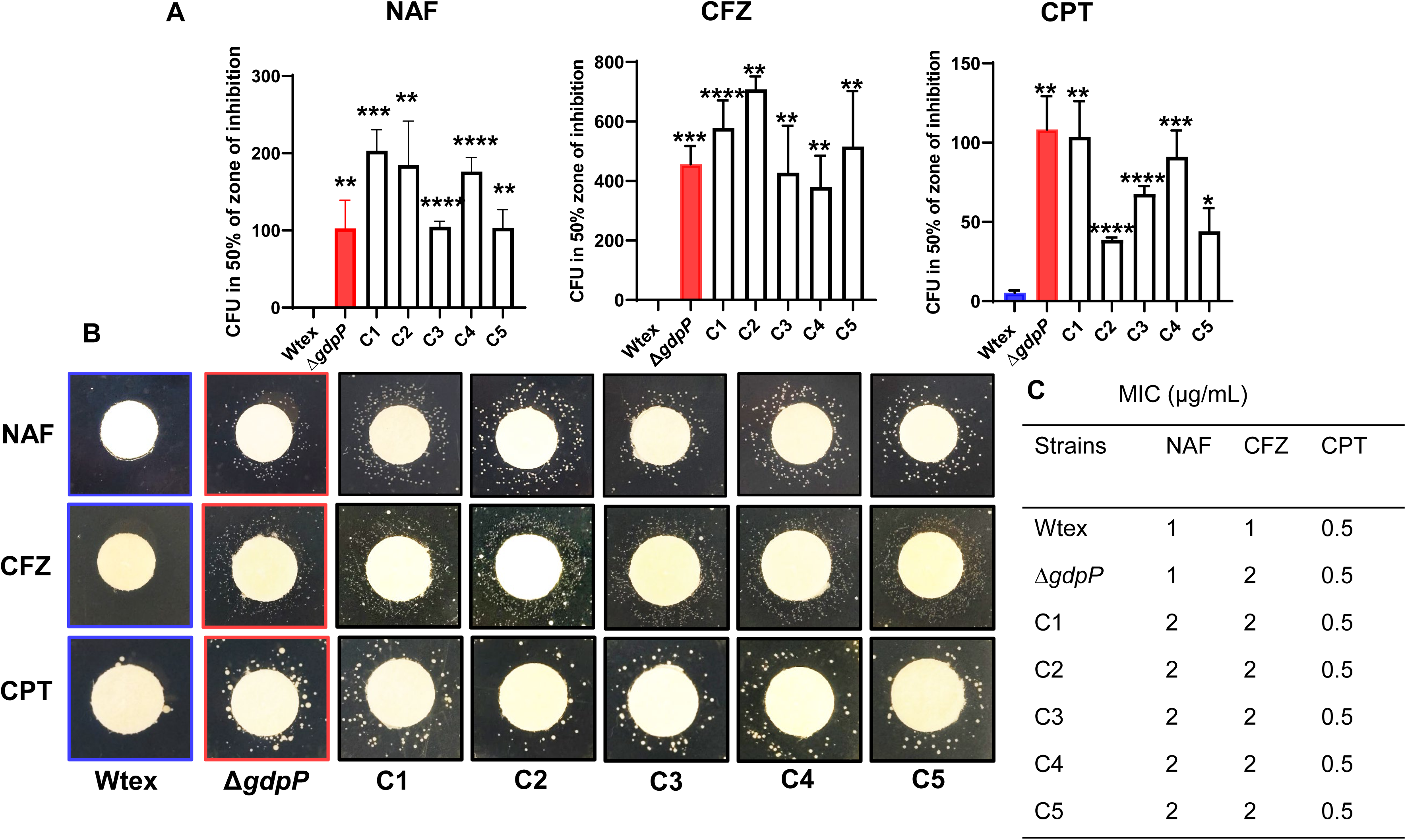
β-Lactam-tolerant Δ*gdpP* colony isolates are not antibiotic-resistant variants. (A-B) Five nafcillin-tolerant Δ*gdpP* colonies (C1 to C5) were purified and analyzed using TD test for their ability to produce tolerance against nafcillin, cefazolin, and ceftaroline. Similar levels of antibiotic tolerance of these isolates to that of their parental strains indicated that the isolated tolerant colonies are not antibiotic-resistant isolates of their parent strain. (C) MICs for the tolerant colony isolates compared to their parental strain. Statistical significance was evaluated using an unpaired t-test. * represents *P* ≤ 0.05, and ** represents *P* ≤ 0.01, *** represents *P* ≤ 0.001 and **** represents *P* ≤ 0.0001.

### Application of bacitracin, a cell wall biosynthesis inhibitor, impairs Δ*gdpP*-mediated β-lactam tolerance in a concentration-dependent manner

Bacitracin binds to undecaprenyl pyrophosphate and inhibits cell wall synthesis by preventing the recycling of the lipid carrier required for the incorporation of lipid II (54) (**Fig.1**). Since bacitracin inhibits cell wall synthesis, we sought to determine its effect on Δ*gdpP*-mediated β-lactam tolerance. We carried out a modified TD test where bacitracin was added in increasing concentrations with a constant dose of β-lactams, nafcillin, and ceftaroline (**Table S3**). We observed a bacitracin dose-dependent reduction in nafcillin and ceftaroline tolerance in Δ*gdpP* (**Fig. 7**). At the highest concentration, bacitracin was even observed to nullify the nafcillin tolerance completely (**Fig. 7A**) and significantly lowered ceftaroline tolerance in Wtex (**Fig. 7B**). Daptomycin, a calcium-dependent cell membrane disruptor, was also tested similarly. Combinational treatment of daptomycin and β-lactams did not affect Δ*gdpP*-mediated tolerance to nafcillin and ceftaroline (**Fig. S3**). These results indicated that bacitracin could potentially impair Δ*gdpP*-mediated β-lactam tolerance in *S. aureus*.

**Figure 7:**
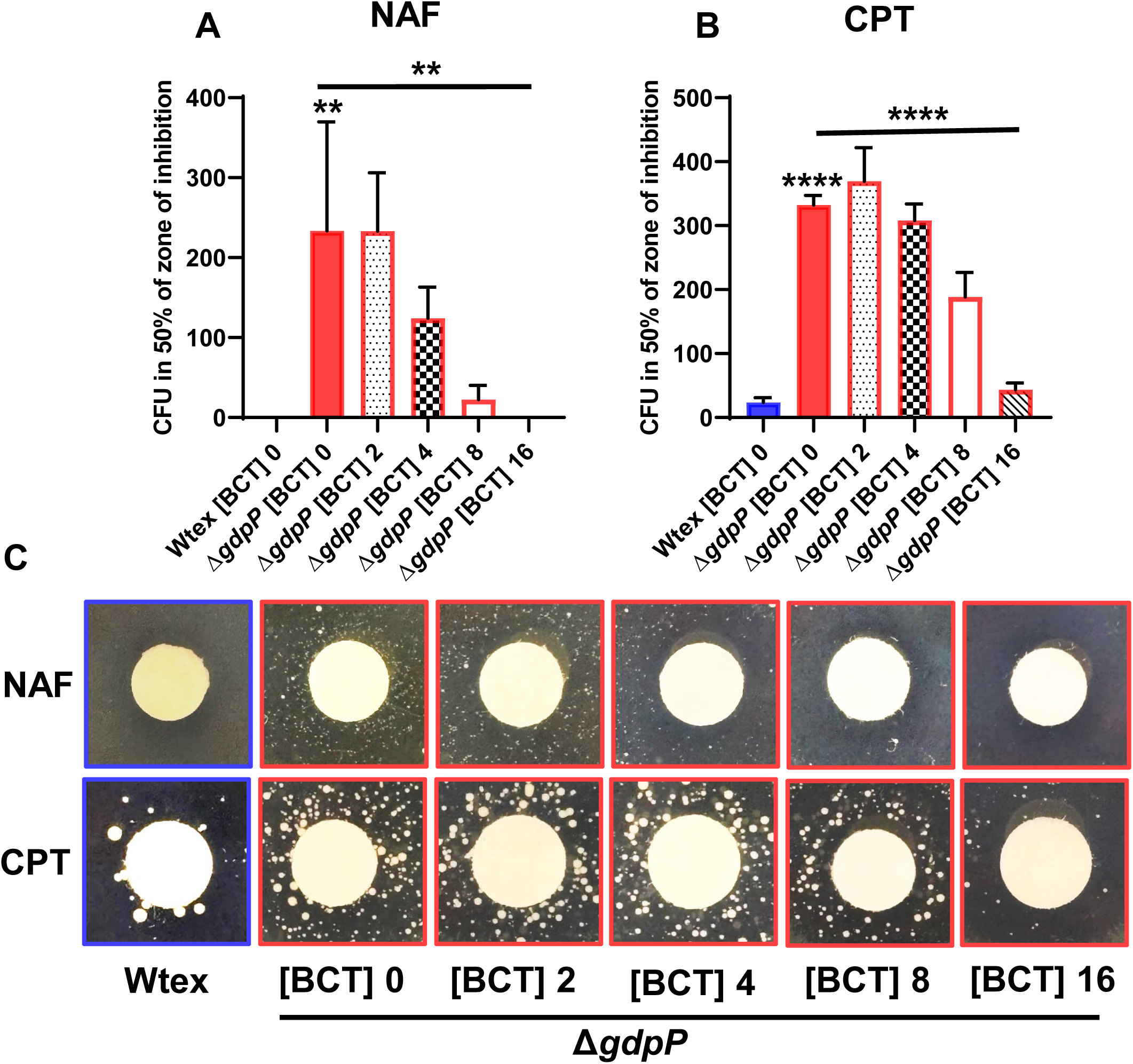
The cell wall synthesis inhibitor, bacitracin, impairs Δ*gdpP*-mediated β-lactam tolerance in a concentration-dependent manner. TD tests were performed using nafcillin (A) and ceftaroline (B), each supplemented with increasing concentrations of bacitracin, indicated with absolute amounts added on the disk in μg. Tolerance to β-lactams significantly decreased with increasing bacitracin levels. (C) Bacterial CFUs observed in the resultant TD tests are shown. Statistical significance was determined using an unpaired t-test. The concentration-dependent decrease in tolerance levels was analyzed using One-way ANOVA. ** represents *P* ≤ 0.01 and **** represents *P* ≤ 0.0001.

### Δ*gdpP*-mediated β-lactam tolerance leads to increased mortality in the *Galleria mellonella* infection model

The *Galleria mellonella* larval model has emerged as a convenient *in vivo* platform to study bacterial infection (55). To determine larval susceptibility to bacterial infection, we inoculated them with Wt, Wtex, and their isogenic Δ*gdpP* mutants, then monitored worm survival with and without β-lactam treatment for 168 h. The control groups, which included heat-killed Wt and nafcillin alone, showed no mortality or melanization in the worms (**Fig S4**). In contrast, infected larvae exhibited mortality beginning at 24 h post-infection, with mortality gradually increasing at subsequent time points (**Fig. 8 and S4**). Progressive melanization, from localized black spots to complete darkening, visually indicated the severity of infection. This phenotypic response flagged larval stress and immune response activation, aligning with previous reports in which healthy larvae remain pale cream, while infection with methicillin-resistant *S. aureus* triggers melanization (**Fig. S4**)

**Figure 8:**
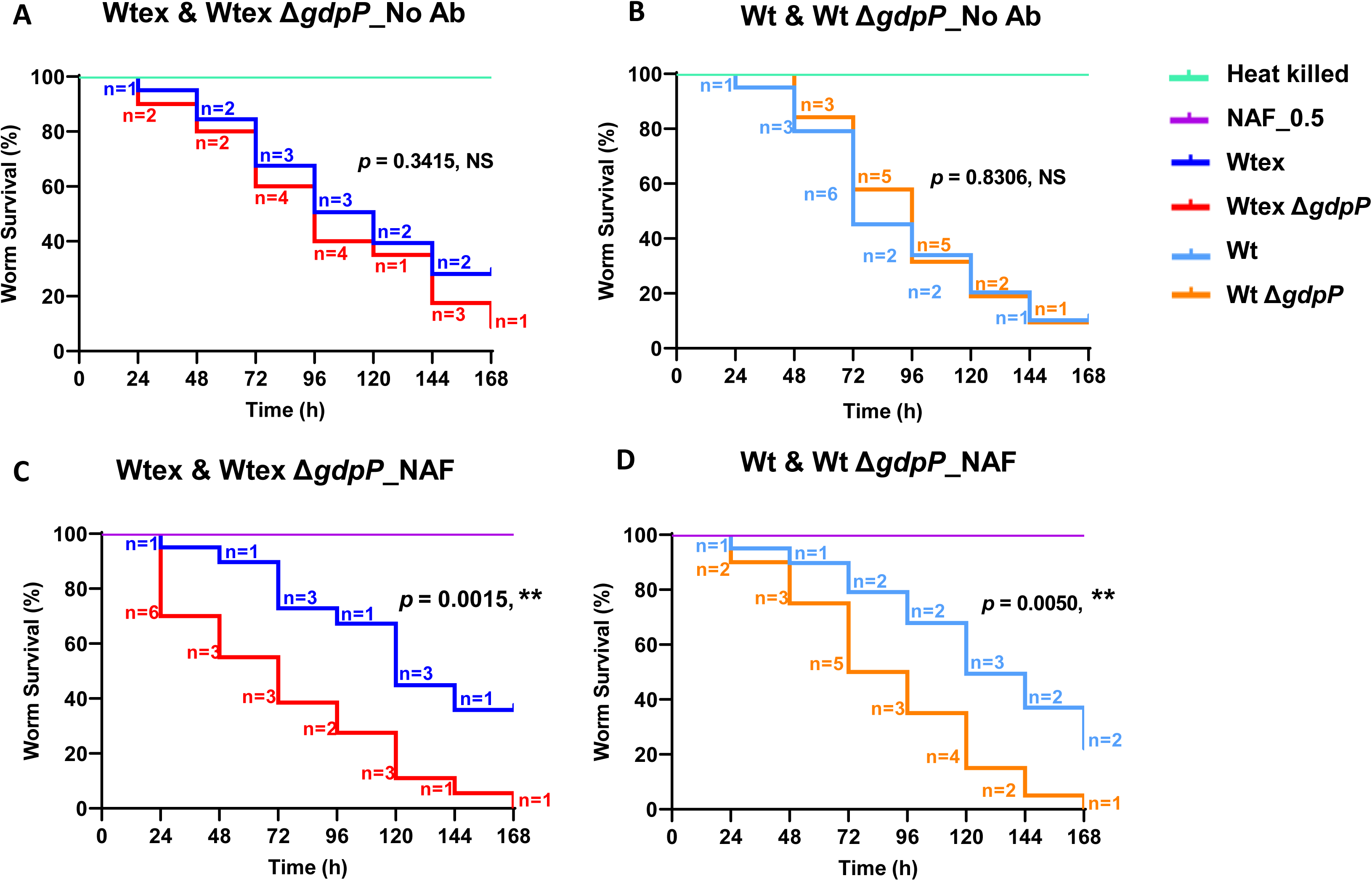
Loss of GdpP function demonstrates β-lactam treatment failure in a *Galleria mellonella* infection model. Kaplan–Meier larval survival plots upon infection with (A) Wtex and Wtex Δ*gdpP* (B) Wt and Wt Δ*gdpP*; without antibiotic treatment. Larval survival under infection followed by subsequent exposure to nafcillin (0.5 µg/mL) treatment 2 h post-infection. (C) Wtex and Wtex Δ*gdpP* (D) Wt and Wt Δ*gdpP*; with β-lactam treatment. Representative images taken at the 168 h time point (**Fig. S4**). Each larvae weighing 200-300 mg were inoculated with 3 × 10⁵ CFU. Heat-killed bacteria and 0.5 µg/mL nafcillin alone are represented as controls. Survival differences among the infection groups were examined with the log-rank (Mantel–Cox) test. Statistical significance ** represents *P* ≤ 0.01 and NS represents non-significant.

(56). In the absence of nafcillin treatment, both Δ*gdpP* mutants exhibited mortality rates higher than those of their Wt strains, although the differences were not statistically significant (**Fig 8A, B**). To evaluate the role of Δ*gdpP-*mediated β-lactam tolerance on worm survival, infected larvae were subsequently treated with nafcillin. Consistent with the results obtained through *in vitro* studies presented above, the Δ*gdpP* mutants exhibited significantly elevated worm mortality during antibiotic exposure, whereas wild-type populations showed noticeable recovery, as evidenced by lower death rates relative to untreated controls (**Fig 8C, D**). These results highlighted that GdpP loss-of-function-mediated tolerance could lead to β-lactam treatment failure during infection.

### Loss of GdpP function mutations promotes persistent invasive infections

We explored within-host emergence of GdpP protein variants among 288 episodes of infection and 663 episodes of nose colonization from CAMERA-2 clinical trial (57). We identified five independent acquisitions of *gdpP* truncations, all during bacteremia (80% MRSA) and after exposure to β-lactams (**Table S4**). This 9-fold enrichment of mutations compared to the background mutation rate was significant (*p*=0.0028, Poisson regression). By contrast, no GdpP protein truncation variants were observed in nasal carriage episodes. Thus, there is a significant selective pressure promoting loss-of-function mutations in persistent bacteremia (adjusted dN/dS 6), while *gdpP* was systematically conserved in nasal carriage (adjusted dN/dS 0.5) (**Fig 9, Table S4**). All bacteremic GdpP mutants emerged under antibiotic exposure, including β-lactams (80%), vancomycin (80%), and daptomycin (40%) (**Table S4**). Interestingly, there was within-host convergent evolution in one case of MRSA bacteremia (**Fig S5**), with two independent mutations during the same episode. Within-host convergence has been described in chronic settings such as *Mycobacterium abscessus* carriage in cystic fibrosis and is thought to represent strong evidence of selective pressure (58) in within-host evolution studies.

**Figure 9:**
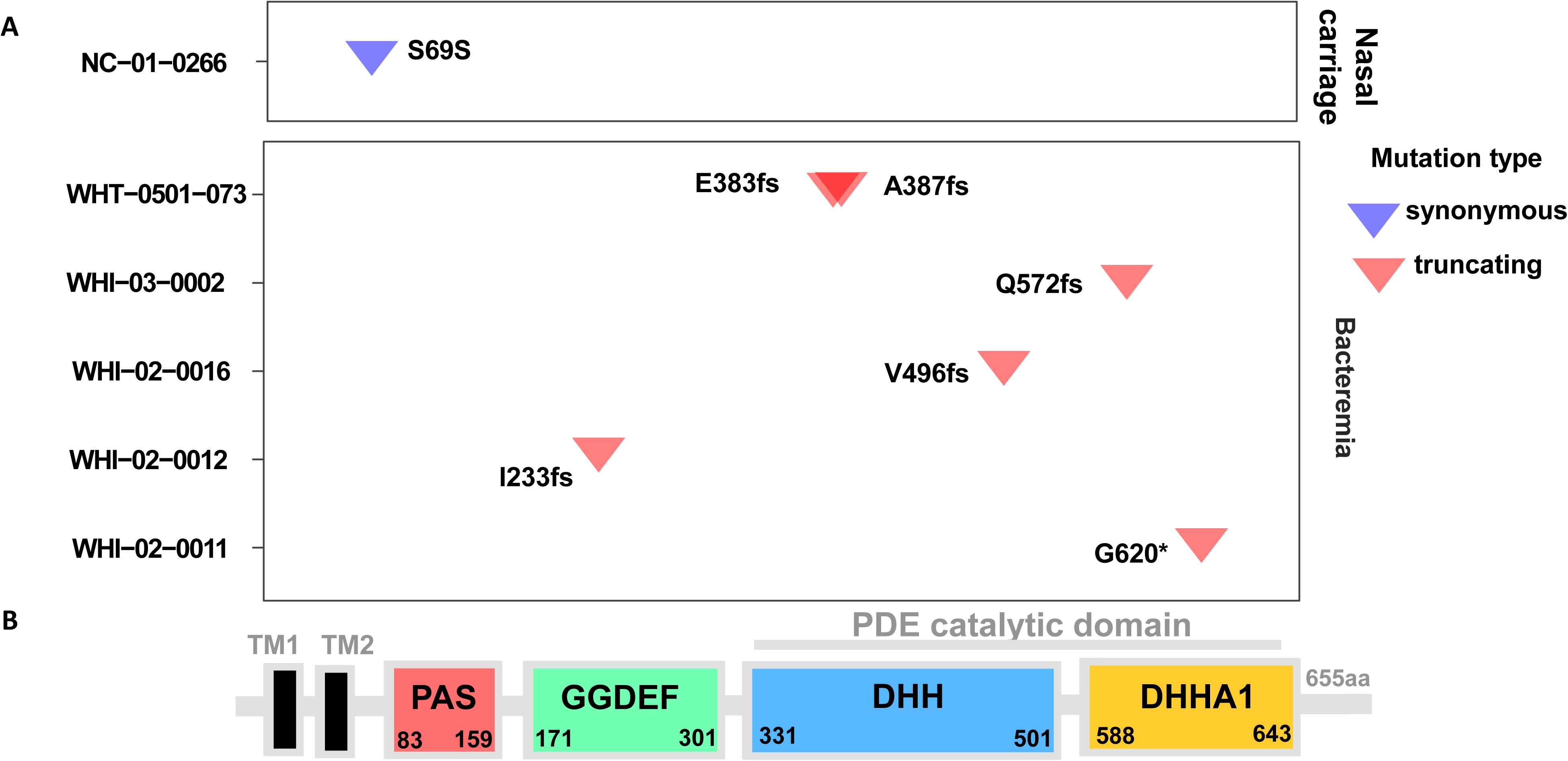
Independently acquired within-host GdpP mutations in publicly available bacteremia and nasal carriage episodes. (A) Map of within-host acquired gdpP mutations in 5 episodes of bacteremia and 1 episode of nasal colonization. Each row represents an infection/colonization episode; mutations are depicted as inverted triangles, colored according to the inferred impact on the gdpP protein sequence. In one episode of bacteremia, a frameshift at position 387 was followed by a second frameshift at position 383. (B) Domain architecture of the gdpP gene, including two transmembrane helices (TM, residues 9-29, 35-54), a degenerate PAS domain (residues 83–159), a GGDEF domain (residues 171–301), a DHH domain (residues 331–501), and a DHHA1 domain (residues 588–643) (adapted from (40)).

## Discussion

*S. aureus* strains are primarily evaluated by testing for antibiotic resistance phenotypes or markers in clinical settings. Antibiotic tolerance is a transient stress response to antibiotics that enables bacteria to survive high antibiotic concentrations through physiological changes without necessarily acquiring resistance-conferring mutations/genes (11) and is challenging to detect in the routine diagnostic environment. This is because the classical MIC metric for resistance measurement is insufficient to detect antibiotic-tolerant bacterial strains. Antibiotic tolerance causes bacteria to become more resilient to antibiotics and promotes faster evolution of antibiotic resistance (11, 59, 60).

Our previous report suggested that *S. aureus* accumulated GdpP loss-of-function mutations at a very early stage during serial passaging experiments carried out in the presence of β-lactam drugs. These mutations led to increased CDA in the bacteria and promoted β-lactam tolerance (14). In addition, several studies have indicated that CDA pleiotropically regulates a broad range of physiological functions in bacteria, including metabolism, genome integrity, virulence, turgor pressure, cell size and volume, osmotic and cell wall homeostasis (16, 24–29, 31, 61–64). The literature also points towards a potentially strong interplay between the multi-level regulatory functions of CDA and drug sensitivity and antibiotic tolerance in multiple bacterial species (28). It has previously been shown that increased levels of CDA mediate tolerance to ceftaroline, a fifth-generation cephalosporin (15, 65, 66). In addition, CDA has been associated with multidrug tolerance, including tolerance to the fluoroquinolone antibiotic ciprofloxacin, sensitivity to antifolate antibiotics and aminoglycosides, and tolerance to vancomycin (17, 65, 67–69). Many of these studies were carried out with either passaged strains or clinically isolated, resistant/tolerant strains. Since these bacterial isolates likely had additional mutations beyond the GdpP loss-of-function mutation, it is harder to determine whether the previously described CDA-related antibiotic-tolerance/sensitivity-associated phenotypes were due to GdpP loss-of-function mutation or to other second-site mutations.

This study offers a detailed insight into which class of antibiotics is most responsible for driving CDA-mediated tolerance in *S. aureus*. The present work establishes several key findings, such as: A) High CDA levels confer tolerance specifically to the β-lactam class of drugs, also including ceftaroline, a highly advanced β-lactam, which is currently used to treat complicated infections caused by *S. aureus* (**Fig. 2-4**). B) GdpP loss-of-function mutations, which are frequently detected among clinical strains (40), confer β-lactam tolerance in both *mecA*+ve (MRSA) and *mecA*-ve (MSSA) background strains (**Fig. S1**). C) β-lactam tolerance positively correlates with CDA concentration in bacteria, suggesting that *gdpP* mutations with varying degrees of loss of function would produce different levels of tolerance (**Fig 5**). D) CDA-mediated β-lactam tolerance can be sensitized by bacitracin, indicating that disruption of lipid carrier recycling and the resulting altered cell wall turnover can counteract the tolerance. E) Δ*gdpP* does not mediate tolerance to vancomycin or daptomycin (**Fig 3, S1**). Prior reports of an association between GdpP loss-of-function mutations and tolerance to vancomycin and daptomycin are likely attributable to secondary mutations present in those isolates (17, 18). It has recently been reported that daptomycin-resistant MRSA have lower levels of cyclic dinucleotides, which supports our results (70). F) Our results from the *G. mellonella* infection model showed that the Δ*gdpP* strain promoted increased worm mortality compared to the Wt, when challenged with a β-lactam. This suggests that GdpP loss-of-function mutations can cause β-lactam treatment failure (**Fig. 8**). G) Gene enrichment analysis showed that loss of GdpP function emerges in persistent invasive infection in both MRSA (*mecA*+ve) and MSSA (*mecA*-ve) strains, suggesting that these loss-of-function mutations might promote treatment failure. Strikingly, neither truncations nor substitutions emerged in nasal colonization, indicating strong purifying selection in the colonization niche and underscoring the importance of a conserved, functioning GdpP for *S. aureus*. Furthermore, a strong association between β-lactam exposure in patients and the detection of the GdpP truncation mutations (**Table S4**) indicates that β-lactam exposure might promote acquisition of such mutations in *S. aureus*.

The exact mechanism of β-lactam tolerance remains unclear. A recent report suggested that in Gram-positive bacteria, CDA-mediated β-lactam tolerance likely involves inactivation of potassium importers such as KupB and the glutamine importer GlnPQ, leading to osmolyte-mediated turgor pressure compensation (71) and activation of the cell wall stress response regulon, the VraTSR system (15). Another recent study has indicated that cell wall stress leads to *gdpP* mutations as part of the stress response. A cyclase regulator, CdaR was reported to sense cell wall defects to activate CDA overproduction (29). The reasons why clinical strains of *S. aureus* tend to accumulate *gdpP* mutations, which are, in most cases, loss-of-function, remain unknown (14, 40). While β-lactam exposure is clearly a primary selective force, host environmental factors may also contribute to the emergence and maintenance of *gdpP* mutations. Mutations in the DHHA1 catalytic domain of GdpP were found in healthcare-associated MRSA strains studied in device-related infections in immunocompromised patients (31). High levels of CDA in bacteria have also been implicated in inducing host immune responses by stimulating the IFN response, the NF-κB pathway, the cGAS-STING pathway, autophagy, and the inflammasome, and by promoting cytokine secretion (70, 72).

Furthermore, there remains a contradiction regarding whether GdpP loss-of-function mutations result in β-lactam tolerance, as reported in this study, or can also confer β-lactam resistance (34, 73). This apparent discrepancy may reflect differences in the genetic backgrounds of the strains examined, as well as the presence of secondary mutations that could act synergistically with GdpP loss-of-function to generate a resistant phenotype. Resolving this issue will require a dedicated future study involving a large and genetically diverse cohort of strains, enabling a systematic assessment of the contribution of genetic background and epistatic interactions to the phenotypic consequences of GdpP inactivation. In summary, our study highlights that enhanced CDA accumulation due to GdpP loss-of-function produces a β-lactam-specific antibiotic tolerance phenotype in *S. aureus*. CDA-mediated β-lactam tolerance could potentially be sensitized through a combinatorial treatment approach with an efficacious antibiotic that acts similarly to that of bacitracin.

## Acknowledgements

Research reported in this publication was supported by the National Institute of Allergy and Infectious Diseases of the National Institutes of Health under award numbers AI165510 and AI195530 to SSC. In addition, SSC would like to thank the Albrecht Foundation for supporting V.D.H. with a 1-year graduate student fellowship. The findings and conclusions presented in this paper are those of the authors and do not necessarily reflect the views of the NIH or the U.S. Department of Health and Human Services. We acknowledge Dr. Joshua Davis and the CAMERA2 study group for providing us with relevant information.

